# Simulating retinal encoding: factors influencing Vernier acuity

**DOI:** 10.1101/109405

**Authors:** Haomiao Jiang, Nicolas Cottaris, James Golden, David Brainard, Joyce E. Farrell, Brian A. Wandell

**Affiliations:** Stanford Center for Image Systems Engineering, Stanford, CA; Psychology Department, University of Pennsylvania

## Abstract

Humans resolve the spatial alignment between two visual stimuli at a resolution that is substantially finer than the spacing between the foveal cones. In this paper, we analyze the factors that limit the information at the cone photoreceptors that is available to make these acuity judgments (Vernier acuity). We use open-source software, ISETBIO^1^ to quantify the stimulus and encoding stages in the front-end of the human visual system, starting with a description of the stimulus spectral radiance and a computational model that includes the physiological optics, inert ocular pigments, eye movements, photoreceptor sampling and absorptions. The simulations suggest that the visual system extracts the information available within the spatiotemporal pattern of photoreceptor absorptions within a small spatial (0.12 deg) and temporal (200 ms) regime. At typical display luminance levels, the variance arising from the Poisson absorptions and small eye movements (tremors and microsaccades) both appear to be critical limiting factors for Vernier acuity.

## Introduction

Many aspects of imaging systems including displays and image coding strategies are designed to accommodate the spatial, chromatic and temporal resolution requirements of the human visual system. Human spatial resolution can be assessed using a variety of experimental protocols (e.g., contrast sensitivity, Vernier acuity, crowding), and for each protocol the bottleneck may be traced to one or several components of the visual system. We used the Image Systems Engineering Toolbox for Biology (ISETBIO) to understand the role that different front-end components play in limiting judgments of Vernier acuity (relative position).

Relative position resolution can be measured by having a subject judge whether a pair of line segments, presented just above and below fixation, is aligned or misaligned. The resolution of this Vernier acuity (relative position) is very precise compared to the sampling rate of the cone receptor mosaic [1] [2]. When the lines are presented to the same eye and near the fovea, observers detect an offset between the lines that is on the order of one-fifth of the width of a single cone. When the two lines are presented in corresponding locations to the right and left eye, the stimulus appears to be a single line whose apparent distance varies with the offset. Discrimination of the binocular offset in this case is called stereoacuity and the threshold is on the order of one third the width of a single cone [3]. The high relative position acuity of the human visual system is a factor driving the need for higher pixel counts in visual displays.

The information used to judge relative position in the Vernier acuity task must be present in the spatial-temporal distribution of the cone absorptions of the human retina, since subsequent processing cannot add to this information. The nature of this information has been understood qualitatively for many years [4]. A thorough analytical assessment of the information available to an ideal observer with a fixed eye position and the signal-defined-exactly and signal-defined-statistically was developed by [5].

This paper extends that analytical work by using computational tools to simulate the encoding and to assess the discriminability of aligned from offset stimuli. The value of creating a computational method is that we can simulate a variety of factors that have no closed-form solution and thus analyze how different biological properties (e.g., physiological optics, eye movements, cone spacing) impact the information encoded for a range of stimuli (chromatic, stimulus size and timing). Here we simulate how Vernier acuity depends upon several specific parameters, including stimulus size, eye movements, image radiance level and defocus.

## Computational methods

### Visual simulation

We refer to the simulation pipeline and inference engine as the Computational Observer (Figure 1) [6]. The simulation specifies quantitatively how the stimulus is transformed to produce the cone absorptions^2^. We use simple machine-learning methods as the inference engine to assess how well different stimuli can be discriminated.

**Figure 1.**
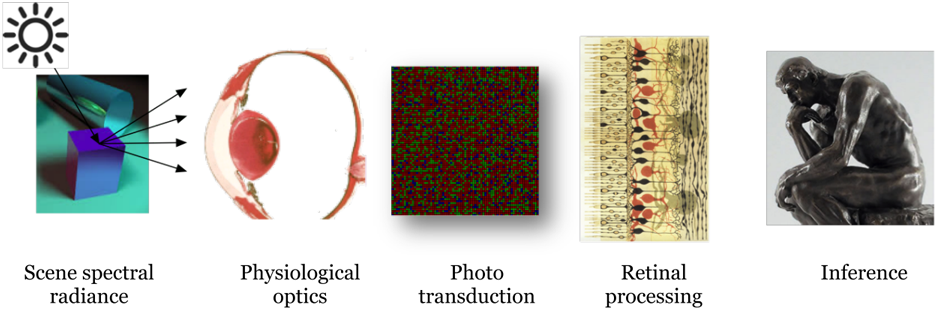
The ISETBIO computational observer for threshold discrimination. We simulate the impact of stimuli at multiple stages within the visual pathway. The stimuli are represented by their spectral radiance, and we calculate the transformation to retinal irradiance by the physiological optics, transmission through various inert pigments, eye movements, and photopigment isomerizations. The software includes methods to calculate photocurrent, bipolar responses and ganglion cell responses. In this paper we focus on the information contained in the isomerizations, a space-time pattern includes the effects of random eye movements and Poisson photon absorptions.

#### Scene modeling

In many psychophysical and engineering problems the stimuli can be described as the spectral radiance from a calibrated display (i.e., a flat surface of Lambertian emitters at a single distance). To fully determine the spectral and spatial / temporal properties of the stimuli, a carefully calibrated display model is required. Calibration involves measurement of the display’s spectral, spatial and temporal properties [7] [8] [9-11].

ISETBIO can also be combined with quantitative computer graphics using modern ray tracing software [12] [13] that generate a description of the light field incident at the cornea [14]. This capability is not used in this article.

#### Physiological optics

The physiological optics model converts the scene radiance to the retinal irradiance. For the small fields of view analyzed here the calculations are isoplanatic; that is, a shift-invariant, wavelength-dependent, convolution kernel can describe them. The software allows specification of an arbitrary kernel; in this application we parameterize defocus by controlling the Zernike polynomial defocus coefficient [15], which makes it possible to align simulations with empirical measurements using adaptive optics [16].

#### Eye movements

We model three types of fixational eye movements: drift, micro-saccade and tremor as independent factors [17] [18] [19].

#### Cone absorptions

Visual quantum efficiency is the product of the lens and macular pigment transmittances and LMS photopigment absorbances [20]. The several components (e.g., macular pigment, lens pigment and photopigment densities and the spatial arrangement of the cone samples) are parameterized for simple experimentation.

#### Retinal neurons

The software includes simulations of photocurrent, bipolar cells and retinal ganglion cells. We do not use these functions in this paper.

#### Inference engine

We train linear support vector machines (SVMs) [21] [22] to discriminate between the spatiotemporal pattern of cone absorptions generated by pairs of test stimuli. We train on sample data and test using an independent set of stimuli. For efficiency, and without losing significant precision, we reduce the cone absorption patterns in the training and test data sets using principal components. The computational observer’s performance is a lower bound on resolution. Using informal experiments, we observed that different classifiers generate somewhat different absolute performance levels. In all cases we tested, however, the general trends and relative performance are consistent. The performance of the classifiers used here does not differ greatly from the ideal classifier with the signal-defined-exactly [5].

### Classification

Figure 2 summarizes the simulation and classification pipeline. We create the dynamic spectral scene radiance for a pair of stimuli - aligned and offset vertical white lines on a gray background. The temporal sequence is sampled at 10 ms over a 400 ms total stimulus duration. The line stimulus luminance was modulated by a Gaussian temporal window with a 100 ms standard deviation, centered at 200 ms. The scene spectral radiance was modeled using calibration data from a conventional (Apple) LCD.

**Figure 2.**
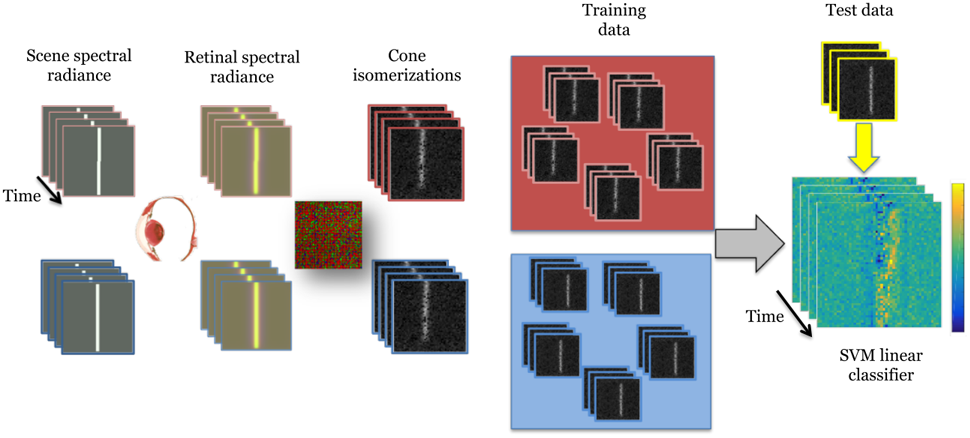
Computational pipeline for Vernier acuity. The Vernier acuity experiment compares two dynamic spectral radiance scenes, one with aligned (blue outline) and one with misaligned (red outline) line segments. We compute the dynamic retinal irradiance and the spatio-temporal patterns of Poisson cone absorptions including eye movements. Multiple samples of the these cone absorptions from the aligned and misaligned stimuli are used to train an SVM linear classifier. Classification accuracy is assessed using independently generated test data (yellow).

We transform the scene spectral radiance into the dynamic retinal spectral irradiance. We then simulate a spatially-regular cone mosaic with randomly interleaved L, M and S cones with 1.4 x 1.4 um apertures (1.96 um^2^), spaced at 2 um, in an L:M:S: ratio of 6:3:1. We calculate the time series of cone absorptions (Poisson noise) for the aligned and offset patterns 1000 times. We then create 1000 eye movement paths in which tremor is sampled at between 76-90 Hz with a mean displacement of 0.4 cones per sample. Drift is assigned a speed of 10 +/- 1 cones/sec, which amounts to a mean drift on the order of 4-6 cones for the 400 ms stimulus duration. The microsaccade frequency is low and rarely occurs during the 200 ms stimulus duration; hence they are immaterial to the simulation. The same eye 1000 movement paths are assigned to aligned and offset data so that small statistical differences in the eye movements cannot contribute to classification performance.

We train the SVM linear classifier on a random selection (80%) of the aligned and misaligned samples. The classifier finds an optimal affine transform that serves as a decision boundary. We estimate classifier accuracy using the held out data (20%). The scripts used to produce the analyses in this paper are available from https://github.com/isetbio/WLVernierAcuity.

## Computational experiments

### Eye movements

The computational methods enable us to explore the interactions between different components of the visual system. Our first experiment considers the impact of two types of eye movements, tremor and drift, on Vernier acuity thresholds (Figure 3). Microsaccades are not important for these experiments with briefly presented stimuli, and there is some reason to believe people suppress microsaccades during these acuity experiments [17],

**Figure 3.**
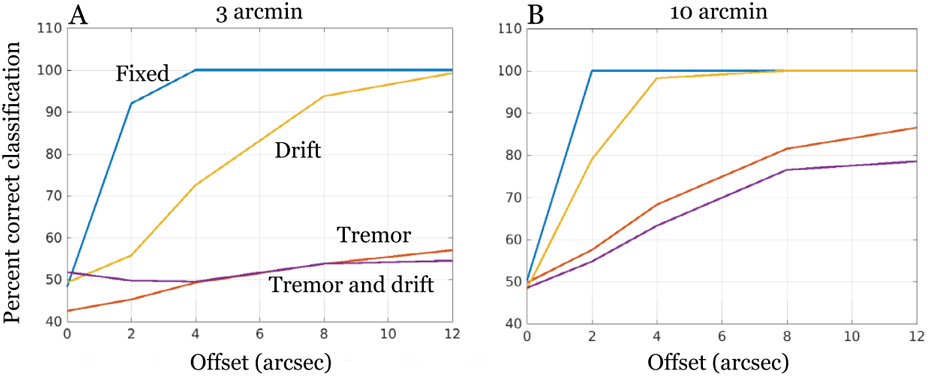
Eye movements impact classification performance. The curves show classification accuracy as a function of bar offset for a different eye movement simulation and bar length. In the absence of eye movements (blue) classification is best, approaching 1 arcsec at 80% correct. In the presence of drift (yellow), tremor (red) or both (purple) threshold declines. The impact of eye movement is particularly large for simulations with a 3 arcmin bar length (A) and the same trends can be observed for simulations with a 10 arcmin bar length (B). Notice that for these two bar lengths when the eye is fixed classification performance is similar, and eye movements have a particularly strong effect on the short bar length. The simulated cone mosaic extended 0.35 deg, cone spacing 2 um, and a random positioning of the L, M, and S cones. The stimuli were white bars on a gray background of 35 cd/m. Stimulus duration was 400 ms with the lines coming on and off with a Gaussian envelope with a standard deviation of 100 ms (effectively a 200 ms stimulus duration).

When the eyes are fixed, as in the signal-defined-exactly case, classification performance is very high (80% correct at 1 arcsec). Introducing drift, tremor or both significantly reduced classification accuracy and, in all cases, the dominant effect is tremor. We performed simulations using a short (3 arcmin) and longer (10 arcmin) line length. The impact of the eye movements is particularly large for the short bar length (Figure 3A). The substantial impact of tremor on the Vernier acuity threshold was a surprise to at least one of the authors. We discuss empirical papers aiming to clarify the impact of eye movements on Vernier acuity [23] [24] later.

### Luminance, defocus, and bar length

Vernier acuity is limited by a combination of Poisson noise in the absorptions, the cone sampling geometry, cone absorption noise arising from random eye movements, and the local loss of signal contrast due to defocus [5]. Increasing stimulus luminance increases the signal-to-noise of cone absorptions; eye movement tremors and defocus both spread the retinal image of the line and decrease the signal-to-noise.

We varied the stimulus radiance level by varying the white point luminance of the simulated display. Vernier acuity increases as display white point luminance increases (Figure 4, left panel). Note that thresholds plateau at 1 arcsec; this may be the threshold level set by eye movements alone.

**Fig 4.**
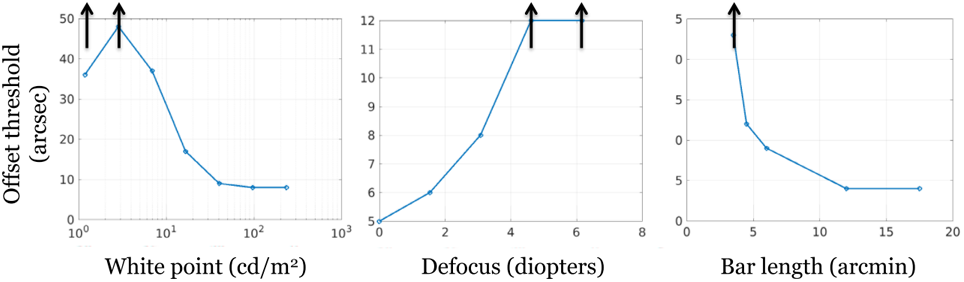
Vernier threshold depends on display luminance, defocus and bar length. We estimated Vernier offset threshold (80% correct) by varying different stimulus or eye model parameters. (A) Threshold varies substantially with the display white point luminance. Threshold decreases as luminance increases, though note the leveling at the highest luminances (B) Threshold increases as we increase optical defocus. The inset images above the graph show the point spread function at 550 nm for the in-focus and 6.15 diopters of defocus. (C) Threshold declines as bar length increases. Arrows indicate points where performance never reached 80%. Stimuli and cone mosaic properties as in Figure 3.

We estimated Vernier offset threshold as a function of defocus (Figure 4, middle panel). Defocus broadens the point spread function, decreases the signal-to-noise at each cone, and increases threshold [5].

Vernier acuity thresholds also decrease as line length increases (Figure 4, right panel). Unlike human performance, the computational observer classification accuracy increases to at least 12 arcmin line [25] [26] [1] [27]. Accuracy in this simulation would have increased further but we only simulated a cone mosaic extending only 0.6 deg (36 arcmin) and in the presence of eye movements a portion of the longer lines sometimes falls beyond the mosaic.

Not shown, we simulated the effect of sweeping out stimulus temporal duration between 50 and 600 ms. We find that given the drift and tremor parameters, threshold decreases up to a stimulus duration of 200 ms.

### Modeling psychophysical measurements

The main analyses explore the experimental and visual system parameters with the aim of dissecting the impact of parameters on limiting resolution. In this section, we coordinate the simulation with specific experimental measurements.

Westheimer and McKee [1] studied Vernier acuity as a function of bar length using bright line targets on a zero background for 200 ms. They report that the line stimulus information is integrated up to about 0.12 deg. We simulated their stimulus conditions and applied the computational observer analysis to a cone mosaic of 0.12 deg (Figure 5). The classification threshold (offset at 75% correct) in the simulation is similar to human performance (Figure 5B).

**Figure 5.**
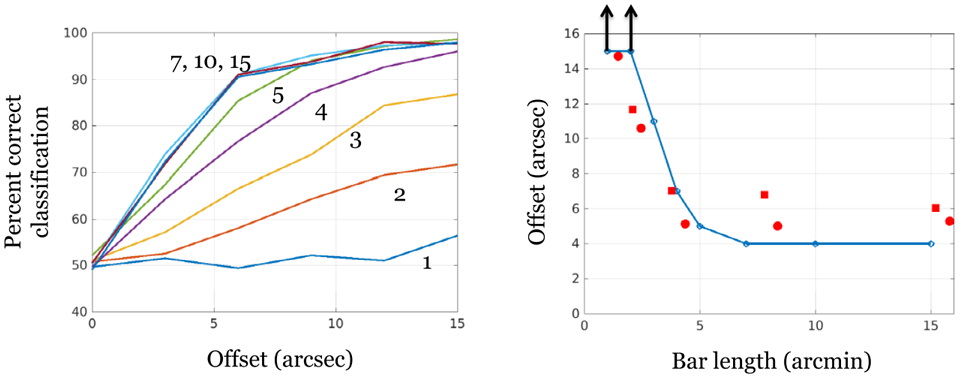
A phenomenological model to summarize performance in Westheimer and McKee (1977). (A) Classification performance is plotted as a function of Vernier line offset for several bar lengths (arcmin). The simulation parameters were set to approximate the subjects described in McKee and Westheimer (1977) - cone mosaic size 0.12 deg, standard eye movements, and a 1.45 arcmin white bar presented for 200 ms on a black background. (B) Vernier offset threshold (80% correct) as a function of bar length is plotted for the SVM linear classifier (blue curve) along with data from two subjects (red symbols). The arrows indicate conditions when the 80% threshold was not quite reached.

The computational observer is intended to characterize an observer making full use of the information available at some location within the visual system, in this case the cone absorptions. In most cases, we expect that additional noise and processing imperfections in other parts of the nervous system will produce behavior that is significantly worse than the computational observer. For Vernier acuity, however, the general agreement between computational observer and human thresholds is surprisingly close. Hence, these analyses support the hypothesis that for the Vernier task the visual system makes efficient use of the information available in the absorptions over a small field of view [5].

## Discussion

### Signal-to-noise in the cone absorptions

Eye movements (Figure 3), luminance level, defocus and bar length (Figure 4) and other factors (e.g., stimulus duration) combine to set limits for Vernier acuity. One way to understand how these factors all contribute is to view how each influences the signal-to-noise in the pattern of cone absorptions (Figure 6). The panels illustrate the effect of defocus that lowers the peak number of absorptions and decreases the slope at the margin of the retinal image of the line. Defocus reduces the ability to detect small differences, and similar curves can be created to show a similar variations in discriminability caused by eye movements and luminance level.

**Figure 6.**
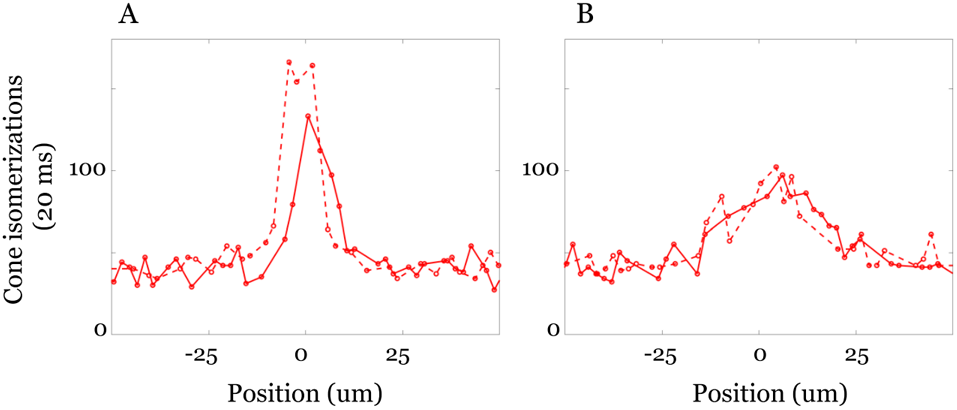
The effect of optical defocus on the spatial pattern of L-cone absorptions. The curves show absorptions in a row of a cone mosaic (2 um cone spacing) with the three types of cones at randomized positions. The absorptions are shown through cones seeing the upper (solid) and lower (dashed) portions of the Vernier lines. The simulation was for a 20 ms stimulus integration time and imaged through (A) average human optics or (B) human optics with a defocus of 2 microns (6.15 diopters). The signal-to-noise and slopes at the edge of the line of the in-focus condition afford more information for judging the alignment.

Vernier acuity threshold depends on bar length [2]. For very short bars, the photons from short bars significantly overlap. Lengthening the bars provides more cone data and also produces absorptions that are clearly assigned to one or the other line [28].

### Related work

The significant impact of eye movements on the performance of the computational observer can be considered in the context of a classic theory of visual acuity: namely, that the dynamics of the signal arising from eye movements is an essential element of visual acuity. This theory has roots going back more than a century, and in certain forms it has been denied [29] [2] [23] [4]. A new form of the theory, connecting acuity with tremors, has been brought forth again in recent work [19] [30].

In the specific case of vernier acuity, empirical measurements show that stabilizing the image [23] or placing it into active motion [24] [27] have very little impact on the threshold. The computational observer experiments show that the presence of small eye movements raises the Vernier threshold. In the fixed eye case the classifier could discriminate as finely as 1 arcsec, while in the presence of eye movements the threshold was raised to 6 arcsec for a 10 arcmin line and much higher for a 3 arcmin line.

These findings raise the question of why there is no difference between the stabilized and unstabilized thresholds measured psychophysically. One possibility is that the visual system anticipates that eye movements will be present. In that case, the neural apparatus needed to resolve the information available without eye movements may not be present. Thus, when eye movements are eliminated by image stabilization, performance does not improve. The situation is analogous to the contrast sensitivity function measured through the S-cones with short wavelength light. Overcoming chromatic aberration and placing a high spatial frequency grating onto the retina does not improve contrast sensitivity; the nervous system is structured under the assumption that chromatic aberration is present and no such retinal images can arise.

Geisler and Davila [28] analyzed the performance limits for an ideal observer case when the signal is defined exactly (SDE) or when the signal is defined statistically (SDS). They estimated thresholds for stimuli comprising dots, similar to the short bar lengths. One might view the contribution here as an extension, some thirty years later, of the computational aspects of that work.

## Conclusion

The computational observer method separates the impact of various factors that comprise a system, whether biological or engineered. The complexity of these factors and their interactions generally preclude the ability to examine their effects by analytic calculations or direct experimental tests. The computational observer method complements the analytical and empirical work.

We see the strengths and weaknesses of analytical theory in many scientific fields. Shannon’s information theory is a fundamental guide to information content, but it is of modest help in understanding why the Internet is slow. The Hodgkin-Huxley equations are compelling, but they do not explain how neuronal signaling depends on the number of myelin wraps and local field potentials. Computational methods provide a useful tool to complement the understanding derived from basic theoretical principles.

This paper reports one of our initial forays using a computational observer approach to understand the visual system. We hope that further development of these models will clarify the interactions between the factors that shape the information available at different stages of the visual encoding. We further hope that experiments and analyses using the computational observer can guide decisions about behavioral and neurophysiological experiments that are needed to test and extend our understanding.

## Acknowledgements

We thank the Simons Foundation Collaboration on the Global Brain for support.

## Footnotes

1 Available at: https://github.com/isetbio

2 Photons can be absorbed by the photopigment without causing a change in the arrangement of the atoms (isomerization). The simulations in ISETBIO do not distinguish between these cases and we simply use the word absorptions.

